# Functional Reorganization of Motor Subcircuits in Parkinson’s disease

**DOI:** 10.64898/2026.02.09.704910

**Authors:** Constantina Theofanopoulou, Neha Bajaj, Alberto Muñoz Sánchez, Bruce Crosson, Steven L Wolf, Venkatagiri Krishnamurthy, Keith M. McGregor, Madeleine E. Hackney

## Abstract

Parkinson’s disease disrupts motor control across multiple body parts, yet the neural mechanisms underlying these impairments remain incompletely defined. We compared resting-state functional connectivity in people with mild-to-moderate Parkinson’s disease (*n* = 58) and neurotypical older adults (*n* = 24), focusing on regions implicated in internally generated (IG) and externally generated (EG) movement pathways. For our analysis, we leveraged the reproducible NeuroMark independent component template and motor effector-specific mapping of primary motor cortex (M1). Our results reveal both increased and decreased connectivity patterns in Parkinson’s disease: M1 subregions associated with control of the leg, hand, and larynx showed robust increases in connectivity exclusively with cerebellar territories, particularly Crus II and Lobules VIIIa/VIIIb. The postcentral gyrus (primary somatosensory cortex) showed primarily increased connectivity with cerebellar regions and the insula. In contrast, the caudate nucleus displayed a mixed profile, with increased connectivity to the superior temporal gyrus and decreased connectivity to the superior medial frontal gyrus and cerebellar Crus II. Our motor effector-specific analysis of disease severity scores (MDS-UPDRS) in people with Parkinson’s disease revealed mild impairments across all categories (leg, hand, larynx) but disproportionately greater hand-related deficits, suggesting that some of the observed M1 connectivity differences may be influenced by these behavioral asymmetries. These anatomically precise, effector-specific alterations suggest compensatory recruitment of cerebellar circuits in Parkinson’s disease and provide a framework for targeting motor subcircuits in rehabilitation, including dance-based interventions.

## Introduction

Parkinson’s disease is the second most prevalent age-associated progressive neurodegenerative disorder.^1^ Nearly 90,000 people with Parkinson’s disease are diagnosed each year in the United States alone,^1^ and over 6 million people live with the disease worldwide.^2^ The incidence of Parkinson’s disease increases markedly with age, affecting an estimated 3% of individuals aged 80 years or older.^3^ Clinically, Parkinson’s disease is defined by four cardinal motor symptoms: rigidity, tremor, bradykinesia, and postural instability.^4^ Rigidity, present in nearly 89% of patients, manifests as an extreme reduction in both upper and lower limb mobility, leading to stiffness and reduced joint range of motion.^5^ Tremor affects approximately 80% of patients, typically presenting as a unilateral resting tremor in the distal extremities.^6^ Bradykinesia, another hallmark feature, involves slowness of movement initiation and execution, and progresses from subtly slowed reaction time and reduced amplitude of limb movements to the loss of spontaneous movements such as arm swing during walking, blinking, or facial expression^6^. Postural instability, characterized by impaired balance and diminished postural reflexes, is most observed in the later stages of Parkinson’s disease and is a major contributor to falls.^7^ Together, these symptoms contribute to significant motor disability in people with Parkinson’s disease.

Although these cardinal symptoms are often discussed in the context of limb motor control, they reflect a broader impairment of motor systems. For example, bradykinesia and rigidity also impair the control of laryngeal muscles, leading to reduced movement amplitude and impaired coordination during phonation, a speech disorder known as hypokinetic dysarthria.^8–10^ This condition is characterized by imprecise consonant articulation, monotony, reduced loudness, altered speech rate, and abnormal pitch.^11–13^ Such impairments reduce speech intelligibility and limit vocal expressiveness, thereby significantly impacting communication and quality of life in people with Parkinson’s disease.^12,14–19^

Previous neuroimaging studies have sought to uncover the neural mechanisms underlying these motor deficits in Parkinson’s disease. Resting-state functional magnetic resonance imaging (rsfMRI) is a particularly powerful approach, as it enables the characterization of spontaneous brain network activity without the performance demands of a task. This method has been widely used to compare functional connectivity patterns in people with Parkinson’s disease, often against control groups such as healthy younger adults^20^ or, more commonly, healthy older adults (HOA).^21^ However, the literature remains far from consensus regarding the specific connectivity alterations in Parkinson’s disease, with discrepancies likely stemming from methodological heterogeneity. Prior studies differ in analytic approach (whole-brain analyses^22^ vs. region-of-interest analyses^23–25^), in the granularity of regions examined (broad networks, such as the dorsal attention network,^23^ vs. specific cortical or subcortical areas^24^), in sample size (e.g., smaller cohorts in Kaut et al., 2020^26^ vs. larger cohorts in Ragothaman et al., 2022^23^), and in interpretive frameworks (e.g., Rodriguez-Sabate et al., 2019^25^; Veréb et al., 2022^22^). This diversity complicates direct comparisons across studies and limits the translational potential of prior findings.

One promising approach to resolving inconsistencies is to frame the analysis within the well-documented distinction between brain pathways supporting internally generated (IG) versus externally generated (EG) movements. IG movements, such as self-paced walking or spontaneous humming, are initiated without direct external cues and rely heavily on cortico-striatal circuits.^27^ EG movements are triggered by external sensory stimuli, e.g., stepping to a metronome or swaying to music,^28–30^ and are thought to engage relatively preserved cortico-cerebellar pathways in early to mid-stage Parkinson’s disease.^27,31–33^ Behavioral evidence supports this dissociation: people with Parkinson’s disease often show greater deficits in IG movements than in EG movements, e.g., walking more smoothly when paced by an auditory cue than when walking at a self-selected pace.^34^ Neuroimaging studies further suggest that people with Parkinson’s disease inadequately recruit cortico-striatal circuits during IG movements and compensate through increased engagement of cortico-cerebellar pathways.^27^ This distinction provides a valuable framework for probing the specific circuit-level changes in Parkinson’s disease, and, in turn, for identifying potential targets for motor rehabilitation strategies that leverage preserved pathways.

In the present study, we used rsfMRI to compare functional connectivity patterns between mild-to-moderate people with Parkinson’s disease and HOA, focusing specifically on brain regions previously implicated in IG or EG movement control. Our design emphasizes anatomical specificity, using well-characterized NeuroMark independent components^35^ (ICs) corresponding to discrete cortical and subcortical regions (e.g., Lobule VIIIb of the cerebellum, dorsomedial precentral gyrus) rather than broad network labels (e.g., “default mode network”). We report connectivity differences as explicit pairwise relationships (e.g., increased connectivity between PreCG/M1 and cerebellar Crus II), enabling direct comparison to task-based and other rsfMRI studies. To contextualize the motor relevance of our findings, we examined the overlap of these ICs with effector-specific regions in the primary motor cortex (M1)—leg, hand, and larynx representations—providing a framework for interpreting connectivity changes in relation to body part-specific impairments. By integrating an IG/EG movement framework with anatomically precise connectivity mapping, our study aims to clarify the functional network alterations in Parkinson’s disease and provide a foundation for targeted interventions, including e.g., dance-based therapies, that could leverage preserved EG pathways to compensate for IG deficits.

## Materials and methods

### Participants and Recruitment Criteria

Seventy-two people with Parkinson’s disease (72) and twenty-four (24) age-matched HOA were recruited through various channels, including the Atlanta Veterans Affairs (VA) Center for Visual and Neurocognitive Rehabilitation (CVNR) registry, the VA Informatics and Computing Infrastructure database, the Michael J. Fox Foundation website, the Movement Disorders Unit at Emory University, newsletters and support groups from Parkinson’s disease organizations, educational events, and word of mouth. Participants were selected based on the following inclusion criteria: all people with Parkinson’s disease had received a clinical diagnosis of Parkinson’s disease by a movement disorders specialist, in accordance with the United Kingdom Parkinson’s disease Society Brain Bank diagnostic criteria.^36^ Participants were required to be at least 40 years old and capable of walking 3 meters or more, with or without assistance. All participants met the criteria for undergoing an fMRI scan, including normal hearing (>40dB pure-tone threshold). Participants were excluded if they scored below 18 on the Montreal Cognitive Assessment (MoCA).^37,38^ Other exclusion criteria included peripheral neuropathy, untreated major depression, a history of stroke, or traumatic brain injury. Depression was assessed using the Beck Depression Inventory-II (BDI-II), and a score of ≥30, indicating severe depression, served as the exclusion threshold.^39^ People with Parkinson’s disease were required to have a unilateral onset of symptoms and demonstrate clear symptomatic improvement with antiparkinsonian medications such as levodopa. People with Parkinson’s disease in Hoehn and Yahr stages I-III were included.^40^ Participants with a tremor score greater than 1 on the Movement Disorders Society Unified Parkinson’s disease Rating Scale (MDS-UPDRS) Part III in a lower limb or with moderate to severe head tremor, were excluded. Finally, all people with Parkinson’s disease participants were tested in the OFF-medication state, defined as being more than 12 hours since their last dose of antiparkinsonian medication.

### Participant Characteristics and Effector-Specific MDS-UPDRS Analysis

For all participants in both the people with Parkinson’s disease and HOA groups, we calculated descriptive statistics for the number of male and female participants, mean age (in years), and mean MoCA scores. For the people with Parkinson’s disease group only, we additionally calculated mean number of years since Parkinson’s disease diagnosis, mean MDS-UPDRS total score, mean MDS-UPDRS-III (Motor Examination) score, and mean scores for three effector-specific categories: hand-related, leg-related, and larynx-related. These effector-specific scores were derived from a subset of MDS-UPDRS items we selected *a priori* based on their relevance to each effector. For each participant, raw scores for the items in each category were summed to produce category totals, which were then normalized by the number of items in that category to yield per-item means. Group-level descriptive statistics were calculated for both the summed and normalized scores. To compare performance across effectors, we conducted a Friedman test on the normalized scores, followed by pairwise Wilcoxon signed-rank tests with Holm correction for multiple comparisons. This analysis provided a standardized framework for quantifying relative impairments across effectors in people with Parkinson’s disease.

### Resting-State Functional Magnetic Resonance Imaging Procedure

Neuroimaging data were collected at the Center for Systems Imaging at Emory University using a 3T Siemens Trio scanner with a Siemens 12-channel head coil. Participants were instructed to lie still with their eyes closed for 9 minutes and 45 seconds, allowing their minds to wander. Foam padding was used to minimize head motion. Following the scan, participants were asked whether they had fallen asleep; if so, the scan was repeated. Resting-state blood oxygen level-dependent (BOLD) fMRI data were acquired using a standard echoplanar imaging (EPI) sequence with an iPAT acceleration factor of 2. The scan parameters were as follows: 55 contiguous 3 mm slices in the axial plane, interleaved slice acquisition, repetition time (TR) = 3000 ms, echo time (TE) = 24 ms, flip angle = 90°, bandwidth = 2632 Hz/pixel, field of view (FOV) = 230 mm, matrix = 76 x 76, and voxel size = 3.0 x 3.0 x 3.0 mm. The first three TRs were discarded to allow for scanner stabilization. An anatomical image was acquired using a high-resolution MPRAGE sequence with 176 contiguous sagittal slices, single-shot acquisition, TR = 2300 ms, TE = 2.89 ms, flip angle = 8°, FOV = 256 mm, matrix = 256 x 256, bandwidth = 140 Hz/pixel, and voxel size = 1.0 x 1.0 x 1.0 mm.

### Resting-State Functional Magnetic Resonance Imaging Data Preprocessing

The imaging data underwent initial quality checks by a single rater, followed by automated quality control using the Magnetic Resonance Imaging Quality Control (MRIQC) software. MRIQC evaluates data quality through various metrics, including signal-to-noise ratio, artifacts, spatial smoothness, and motion-related parameters, to identify potential issues that could affect subsequent analyses. This process flagged one dataset (sub131-ses01) as problematic due to quality control concerns, leading to its exclusion from the analysis. Preprocessing included slice-time correction and motion correction on the functional volumes using SPM12. Physiological noise from pulse and respiration was normalized to standard space. The first 8-10 scans were discarded to mitigate saturation effects. The Retrospective Image Correction (RETROICOR) algorithm was applied to remove physiological noise related to cardiac and respiratory fluctuations.^41^ Cardiac and respiratory functions were monitored using photoplethysmography on the left index finger and a respiratory belt around the chest. BOLD signal fluctuations due to low-frequency cardiac and respiratory waveforms were detrended using established methods.^42^ The corrected data were low-pass filtered (cutoff frequency: 0.1 Hz) to isolate low-frequency resting-state BOLD fluctuations.^43^ EPI images were then spatially normalized to the Montreal Neurological Institute (MNI) 3mm isotropic template using nonlinear registration and smoothed with a 10 mm full-width half maximum (FWHM) Gaussian kernel. Signal intensities for each volume were z-transformed, excluding the first six volumes from the calculation of the mean and standard deviation to avoid pre-steady-state outliers.

### Regions of Interest selected within Internally Generated & Externally Generated Pathways

To refine our regions of interest (ROIs) related to IG and EG movements, a literature review was conducted using NCBI PubMed (Supplementary Table 1). Keywords utilized to find relevant studies included: “rhythmic”, “internally generated”, “externally generated”, “cued movement”, “paced movement”, “fMRI”, “motor”, “movement”, “tapping”, and “drawing”. This analysis was assembled based upon a previous review^44^ and was further structured to include original research studies examining people with Parkinson’s disease, HOA, and healthy adults (HA). For classification, HOA were defined as those aged > 55 years. The aims of this review were to delineate a) potential differences in the brain pathways associated with IG and EG movements, and b) potential differences in the IG and EG pathways between these groups (people with Parkinson’s disease, HOA, and HA). The review focused on identifying consistent findings across studies to strengthen the validity of our ROI selection. Studies were chosen based on their relevance to movement pathways, with particular attention to the activation and deactivation patterns in brain regions involved in IG and/or EG movements.

Within the framework of our study, IG movements refer to those that an individual initiates without external cues or prompts. EG movements, on the other hand, are those in which an individual responds to external cues or stimuli, such as auditory, tactile or visual signals.^45^ Examples of IG movements we encountered in the literature included non-cued tasks such as foot tapping, hand tapping, and dorsoplantar flexion. EG movements encompassed cued tasks, such as auditory or visual cue-driven foot tapping, hand tapping, dynamic tracing, and paced finger tapping. By distinguishing these two movement categories, we aim to examine the neural substrates associated with both voluntary, self-initiated movements and those modulated by external environmental cues.

### Independent Component Analysis

Based on the brain regions identified as relevant to IG and EG movements, we selected those consistently identified as associated with these movement pathways (n=27) as our ROIs for further analyses (Table 1). Since the reviewed studies reported different activity peaks for the identified regions, to ensure the replicability of our findings, we focused on brain components within these regions (e.g., M1, CB) that would be as reproducible as possible. For this purpose, we used the NeuroMark pipeline,^35^ which provides well-established and reproducible independent components (ICs) that serve as priors for analysis across pathologies. To enhance the robustness of our results, we mapped our 27 pre-identified resting-state ROIs to the corresponding regions derived from the NeuroMark pipeline. In some cases, a single ROI corresponded to one distinct NeuroMark IC (e.g., SMA to IC 84), which was used as a prior in our analysis. In other cases, an ROI mapped to multiple replicable ICs from the same broader brain area (e.g., PreCG/M1 mapped to three ICs –IC 2, IC 54, and IC 66– differing along the dorsoventral and mediolateral axes of the gyrus).

**Table 1.**
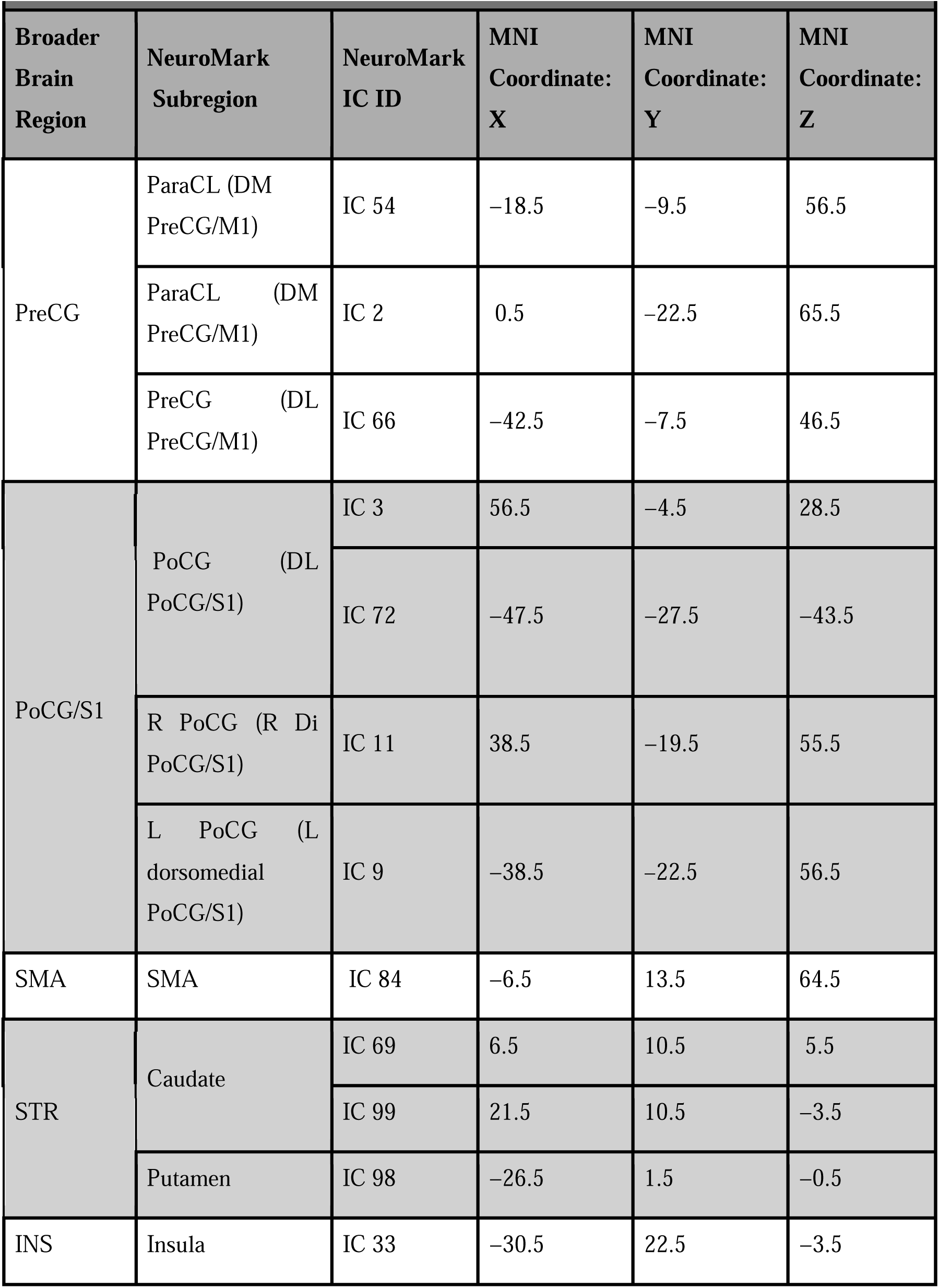
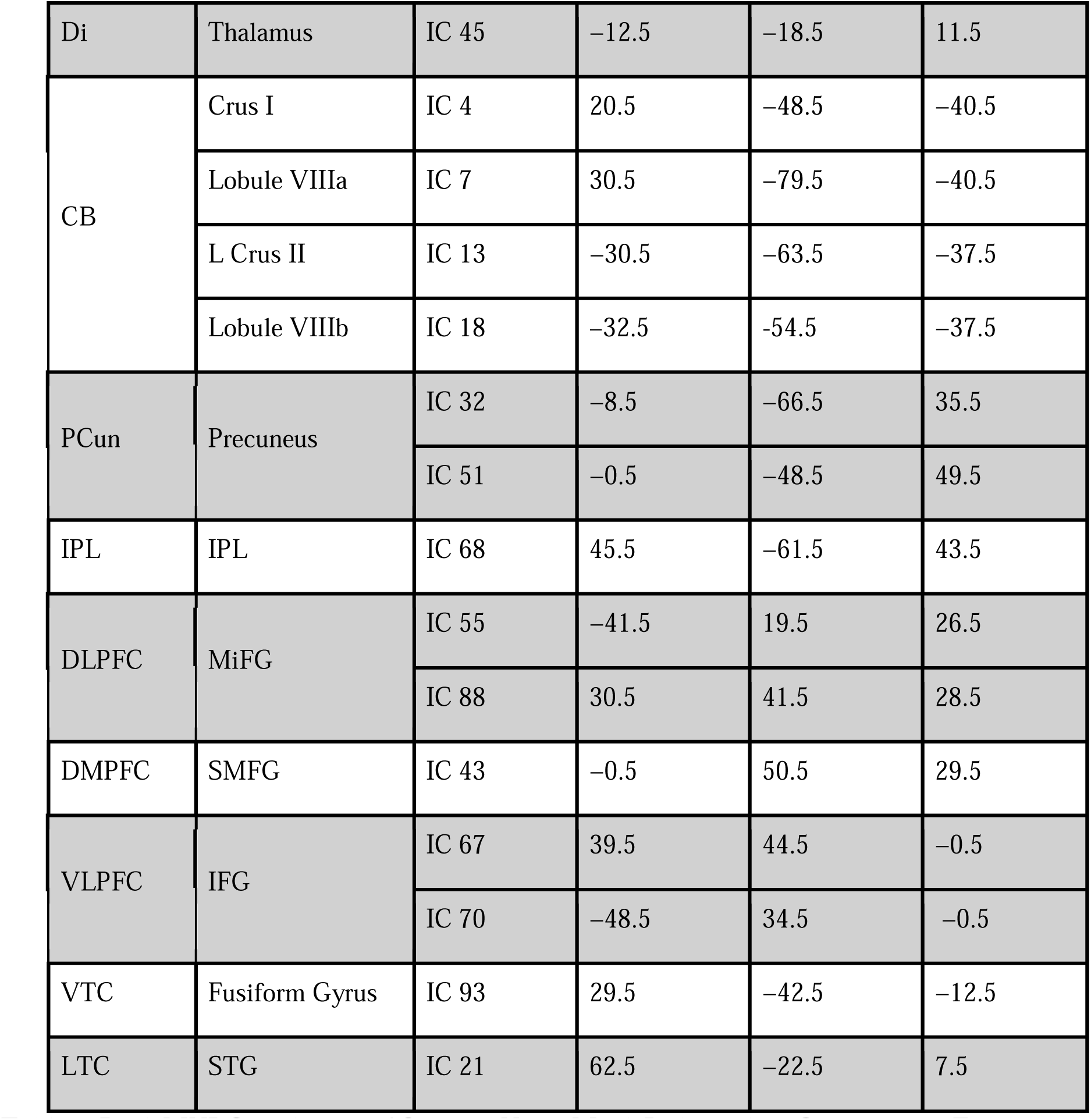
Peak MNI Coordinates of Selected NeuroMark Independent Components. This table shows the 27 NeuroMark ICs selected for analysis, which correspond to the ROIs repeatedly implicated in IG and EG neural pathways in Table 1. Column 1 lists the broader brain region in which each NeuroMark IC is located, aligning with the broader regions outlined in Table 1. Column 2 specifies the subregion corresponding to each NeuroMark component. For regions without specific laterality indicated (i.e., R = Right or L = Left), the priors correspond to both hemispheres. Column 3 provides the index ID of each NeuroMark IC. Columns 4, 5, and 6 present the MNI X, Y, and Z peak coordinates for each NeuroMark component used in the analysis. Broader Brain Region Abbreviations: PreCG; Precentral Gyrus; SMA, Supplementary Motor Cortex; STR, Striatum; INS, Insula; Di, Diencephalon; CB, Cerebellum; PCun, Precuneus; IPL, Inferior Parietal Lobe; DLPFC, Dorsolateral Prefrontal Cortex; DMPFC, Dorsomedial Prefrontal Cortex; VLPFC, Ventrolateral Prefrontal Cortex; VTC, Ventral Temporal Cortex. NeuroMark Subregion Abbreviations: ParaCL, Paracentral Lobule; PreCG, Precentral Gyrus; M1, Primary Motor Cortex; DM, Dorsomedial; DL, Dorsolateral; Di: Dorsal-intermediate; PoCG, Postcentral Gyrus; SMA, Supplementary Motor Area; CB, Cerebellum; IPL, Inferior Parietal Lobe; MiFG, Middle Frontal Gyrus; SMFG, Superior Medial Frontal Gyrus; IFG, Inferior Frontal Gyrus; STG, Superior Temporal Gyrus; LTC, Lateral Temporal Cortex.

Following preprocessing, the resulting time courses were analyzed using the GIFT software package (Group ICA of fMRI Toolbox^46^) to perform spatially constrained independent component analysis (ICA). For each subject and session, the ICA procedure decomposed the fMRI time series into three primary outcome variables: a) component spatial maps: Voxel-wise spatial representations of each independent component, illustrating the extent to which different brain regions contribute to a given network; b) component power spectra: Frequency-domain information for each independent component, describing the spectral power distribution of the associated time series; and c) between-component connectivity (functional network connectivity, FNC): Temporal correlations between the time courses of independent components, offering insights into functional interactions between networks.

### Statistical analysis of functional connectivity

All 27 ROIs, corresponding to NeuroMark ICs, were selected for analysis. Pairwise comparisons of FNC across all ICs were conducted, and the resulting spatial maps, power spectra, and FNC matrices were submitted to group-level statistical analysis using the Multivariate Analysis of Covariance (MANCOVAN) toolbox, implemented in the latest version (as of August 2025) of the Group ICA of the fMRI Toolbox (GIFT). To evaluate group differences, independent t-tests were performed for all pairwise IC combinations. A significance threshold of *P* < 0.01 was applied to identify uncorrected effects. To mitigate false positives from multiple comparisons, false discovery rate (FDR) correction was performed using the MAFDR method (multivariate adaptive false discovery rate^47^), which estimates q-values as the minimum FDR threshold at which a test is considered significant. A threshold of *q* < 0.05 was used to determine FDR-corrected significance.

### Motor Effector Localization within Primary Motor Cortex

Given that several of the Precentral Gyrus/Primary Motor Cortex (PreCG/M1) NeuroMark ICs used in this study were involved in significant rsfMRI differences between people with Parkinson’s disease and HOA, we aimed to determine which motor effectors these components most likely represent. Seminal studies on the functional organization of the M1 have consistently identified three major effector-specific regions: (1) leg, (2) hand, and (3) larynx.^48,49^ To investigate how our ICs may correspond to these subdivisions, we generated activation masks for each effector based on existing literature and compared them to the spatial maps of the NeuroMark ICs.

We conducted a comprehensive literature review using NCBI PubMed, Google Scholar, and JSTOR to identify relevant neuroimaging and brain stimulation studies that reported activation peaks associated with voluntary movement of the leg, hand, or larynx. For the larynx area, we prioritized studies examining M1 activation during speech production, as this effector was considered particularly relevant to the articulation and phonation deficits attested in Parkinson’s disease.^8,50,9,51^ For the hand and leg effectors, we focused on studies that systematically compared activation patterns across multiple effectors or provided robust mappings of effector-specific activity.

Reported peak activation coordinates, either in MNI or Talairach-Tournoux space, were compiled and analyzed using GingerALE, a tool within the BrainMap platform designed to conduct Activation Likelihood Estimation (ALE) meta-analyses.^52^ Coordinates reported in Talairach space were converted to MNI space using the Lancaster transform, as implemented in the Yale BioImage Suite Package. GingerALE aggregates activation peaks across studies to identify regions of consistent activation. ALE maps were thresholded using cluster-level family-wise error (FWE) correction, with a cluster-forming threshold of *P* < 0.01 and a corrected cluster-level threshold of *P* < 0.05. Only clusters showing robust convergence across studies were retained for further analysis. We next visualized both the NeuroMark IC spatial maps and the GingerALE-derived effector-specific masks on a standard template brain using SurfIce. Overlap between ICs and effector-specific regions was reported to help contextualize the potential motor specificity of the connectivity findings.

## Results

### Resting-State Connectivity Differences between People with Parkinson’s disease and HOA

In this study, we investigated resting-state functional MRI (rsfMRI) connectivity differences between people with Parkinson’s disease and HOA, focusing on brain areas previously implicated in IG and EG movement pathways. This approach builds on evidence that internally driven movements, such as spontaneous motor actions, are often impaired in people with Parkinson’s disease, whereas externally cued movements, such as those synchronized to music or rhythm, may remain relatively preserved, especially in earlier stages of the disease.^34^

We first conducted an extensive literature review to identify brain regions associated with IG or EG movements in both people with Parkinson’s disease and neurotypical populations (HOA and HA) (Supplementary Table 1). Regions that appeared repeatedly across studies, such as the Precentral Gyrus (PreCG), Supplementary Motor Area (SMA), Cerebellum (CB), and Striatum (STR), were selected for further analysis (Supplementary Table 1). To enhance reproducibility, we mapped these regions to 27 well-established independent components (ICs) from the NeuroMark pipeline,^35^ which provides validated spatial priors for resting-state networks (Table 1). In some cases, a single ROI (Supplementary Table 1) corresponded to one distinct NeuroMark IC (e.g., SMA to IC 84) (Table 1), which was used as a prior in our analysis. In other cases, an ROI (Supplementary Table 1) mapped to multiple replicable ICs from the same broader brain area (e.g., PreCG/M1 mapped to three ICs –IC 2, IC 54, and IC 66– differing along the dorsoventral and mediolateral axes of the gyrus) (Table 1). In all cases, we included all relevant NeuroMark ICs to ensure comprehensive coverage of each ROI and maximize the inclusion of functionally relevant subregions.

351 pairwise comparisons were performed between the 27 NeuroMark ICs to assess functional connectivity differences between people with Parkinson’s disease and HOA. Independent two-sample t-tests (uncorrected threshold: *P* < 0.01) revealed multiple IC pairs with increased or decreased connectivity in people with Parkinson’s disease relative to HOA (Supplementary Tables 2, 4, and 5). To address the risk of false positives due to multiple comparisons, we applied the Storey^47^ MAFDR correction, with *q* < 0.05 as the corrected significance threshold (Supplementary Tables 3, 4).

Several IC pairs (*n* = 15 pairs) showed increased functional connectivity in people with Parkinson’s disease compared to HOA (*P* < 0.01) (Figure 1, Supplementary Tables 2 and 4). Notably, several of these increases involved ICs from the PreCG/M1, particularly IC2 and IC66. These PreCG/M1 ICs exhibited greater connectivity with multiple cerebellar components, including CB Crus II (IC13), Lobule VIIIa (IC7), and Lobule VIIIb (IC18). For example, dorsomedial PreCG (IC2) displayed elevated connectivity with all three of these cerebellar lobules, while dorsolateral PreCG (IC66) was specifically coupled with CB Crus II and Lobule VIIIb. The postcentral Gyrus (PoCG) ICs also showed increased connectivity with several other areas: the left dorsomedial PoCG (IC9) was more strongly connected with CB regions (Crus I, Crus II, Lobule VIIIb, and right Lobule VIIIa), while all PoCG ICs (IC3, IC9, and IC11, corresponding to right dorsolateral PoCG/S1, left dorsomedial PoCG/S1, and dorsal (intermediate) PoCG/S1, respectively) were found to have higher connectivity with the Insula (IC33). Lastly, the caudate (IC99) also demonstrated increased connectivity with the STG (IC21). Of these pairs, one—IC2 (PreCG/M1) and IC7 (CB right Lobule VIIIa)—survived FDR correction (*q* < 0.05), indicating robust increased connectivity in people with Parkinson’s disease.

**Figure 1.**
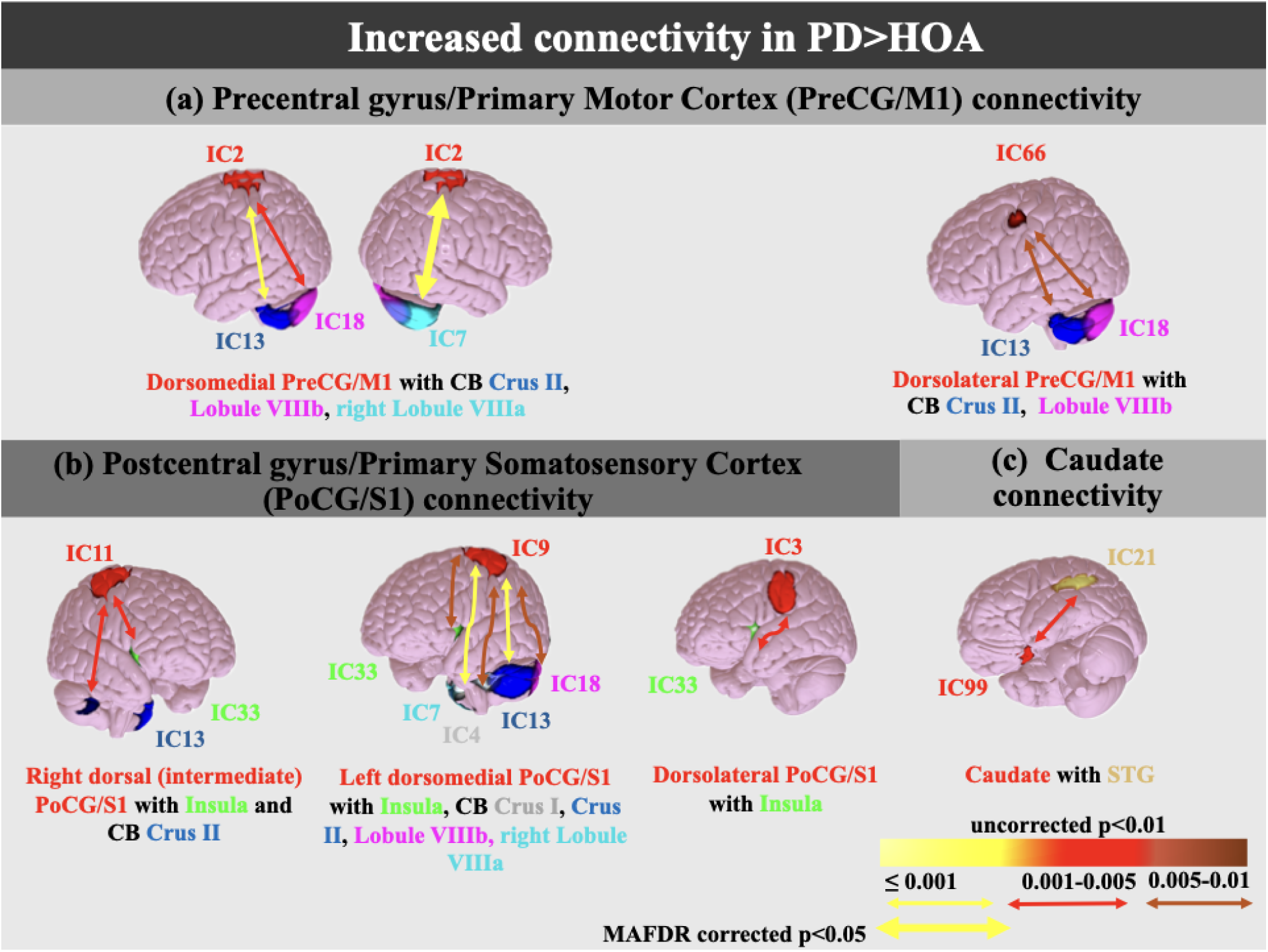
Increased Functional Connectivity in Parkinson’s disease (people with Parkinson’s disease) Compared to Healthy Older Adults (HOA). (A) Precentral Gyrus/Primary Motor Cortex (PreCG/M1) Connectivity. PreCG/M1 ICs—IC2 and IC66—are highlighted in red, with their connectivity targets in other colors. IC2: dorsomedial PreCG/M1; IC66: dorsolateral PreCG/M1 (ventral to IC2). Targets include IC13 (dark blue): Cerebellum (CB) Crus II; IC7 (light blue): right CB Lobule VIIIa; IC18 (purple): CB Lobule VIIIb. **(B) Postcentral Gyrus/Primary Somatosensory Cortex (PoCG/S1) Connectivity.** PoCG/S1 ICs—IC11, IC9, IC3—are shown in red. IC11: right dorsal (intermediate) PoCG/S1. IC9: left dorsomedial PoCG/S1; IC3: dorsolateral PoCG/S1; Targets include CB regions (IC7, IC13, IC18; color coded as in (a)) and IC7 (marble gray): Cerebellum (CB) Crus I, and Insula (IC33, light green). **(C) Caudate Nucleus Connectivity.** IC99 (red): Caudate; Target: IC21 (gold): Superior Temporal Gyrus (STG). Color coding of connectivity arrows reflects significance levels from uncorrected t-tests (*P* < 0.01): yellow (*P* ≤ 0.001), red (*P* = 0.001–0.005), brown (*P* = 0.005–0.01). Bold yellow arrows denote results surviving MAFDR correction (*q* < 0.05).

In contrast, fewer IC pairs (*n* = 4) showed decreased connectivity in people with Parkinson’s disease relative to HOA (*P* < 0.01) (Figure 2, Supplementary Table 2, 3, 5). These included connections between right dorsal PoCG (IC11) and CB Crus II (IC13), and between the right dorsal PoCG (IC11) and left dorsomedial PoCG/S1 (IC9). Decreased connectivity was also observed between the insula and several cerebellar lobules (IC13, IC18, IC7). Lastly, the caudate nucleus showed reduced coupling with the superior medial frontal gyrus (SMFG; IC70) and CB Crus II (IC13). However, none of these reductions survived MAFDR correction.

**Figure 2.**
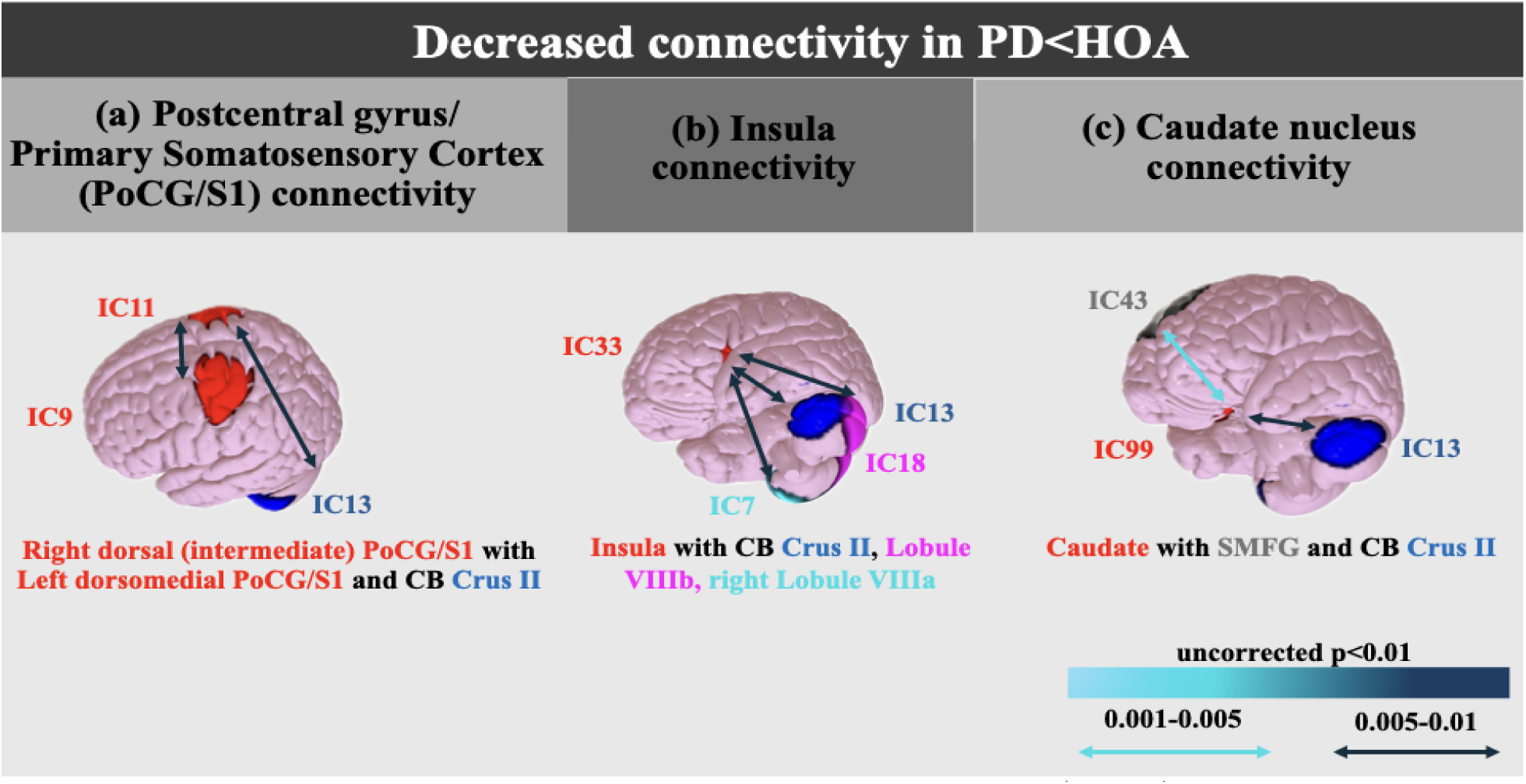
Decreased Functional Connectivity in Parkinson’s disease (People with Parkinson’s disease) Compared to Healthy Older Adults (HOA). **(A) Postcentral Gyrus/Primary Somatosensory Cortex (PoCG/S1) Connectivity**. PoCG/S1 ICs—IC11 and IC9—are shown in red. IC11: right dorsal (intermediate) PoCG/S1; IC9: left dorsomedial PoCG/S1. Targets include IC13 (dark blue): Cerebellum (CB) Crus II. **(B) Insula Connectivity.** Insula IC33 is shown in red. Targets include IC13 (dark blue): CB Crus II; IC7 (light blue): right CB Lobule VIIIa; IC18 (purple): CB Lobule VIIIb. **(C) Caudate Nucleus Connectivity.** Caudate IC99 (red); Targets: IC43 (gray): Superior Medial Frontal Gyrus (SMFG) and IC13 (dark blue): CB Crus II. Connectivity arrow color indicates p-value significance levels from uncorrected t-tests (*P* < 0.01): light blue (*P* = 0.001–0.005), dark blue (*P* = 0.005–0.01).

The data revealed several notable trends. PreCG/M1 components were only associated with increased connectivity in people with Parkinson’s disease, exclusively with CB regions. In contrast, SMFG (IC70) appeared exclusively in decreased connectivity pairs. Other regions, such as the caudate nucleus and PoCG, displayed bidirectional patterns—showing increased connectivity with some regions and decreased with others. For instance, the caudate (IC21) exhibited greater connectivity with the superior temporal gyrus (STG) but reduced connectivity with CB Crus II and SMFG. Overall, these results highlight distinct connectivity patterns across brain regions, with some showing exclusively increased, exclusively decreased, or mixed connectivity differences in people with Parkinson’s disease compared to HOA.

### Motor Effector Localization of PreCG Components

Given the prominence of PreCG/M1 ICs in the observed group differences, we next examined which motor effectors (leg, hand, larynx) these ICs most likely represent. Using coordinate-based meta-analyses (via GingerALE) of studies targeting these effectors, we created effector-specific masks (Supplementary Table 6) and compared them with the spatial extent of PreCG/M1 ICs.^52^ The results (Figure 3) showed that: a) IC2 (dorsomedial PreCG/M1) overlapped primarily with the leg area; b) IC54 (also dorsomedial PreCG/M1, located more ventrally with respect to IC2) overlapped with both the leg and hand areas; and c) IC66 (dorsolateral PreCG/M1, located more ventrally with respect to IC54) showed overlap with the hand and larynx areas. These anatomical distinctions suggest that the observed connectivity increases in people with Parkinson’s disease may reflect effector-specific network differences, such as stronger cerebellar coupling in leg-related (IC2) and hand/larynx-related M1 (IC66) motor circuits (Figures 1 and 3).

**Figure 3.**
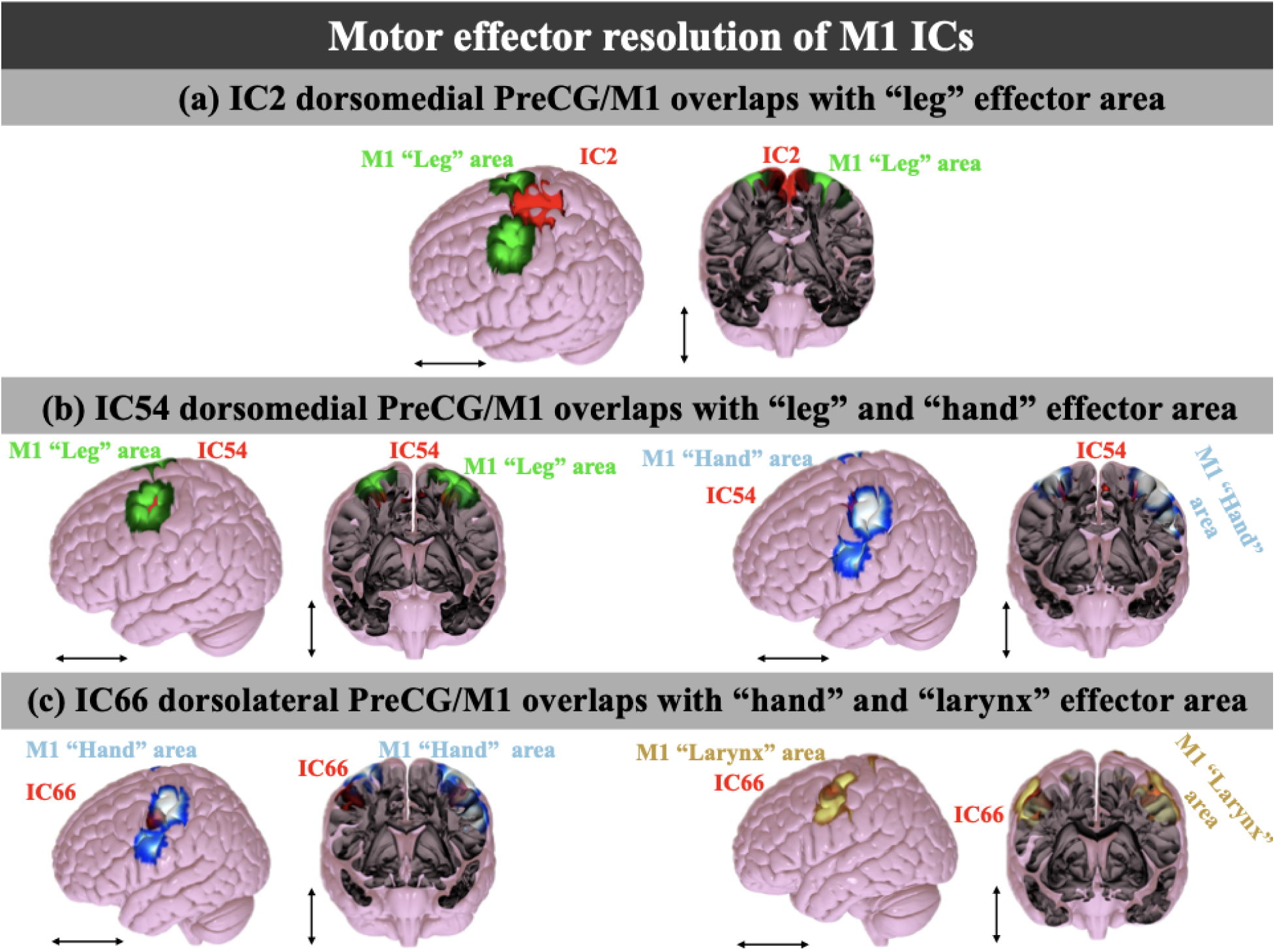
Motor Effector Resolution of PreCG/M1 ICs. ICs are overlaid with GingerALE-generated M1 effector-specific masks. **(A) IC2 (red):** dorsomedial PreCG/M1 overlaps with the leg effector mask (light green). **(B) IC54 (red):** dorsomedial PreCG/M1 (ventral to IC2) overlaps with both leg (light green) (left) and hand (light blue) effector masks (right). **(C) IC66 (red):** dorsolateral PreCG/M1 (ventral to IC54) overlaps with both hand (light blue) and larynx (gold) effector masks. Black arrows indicate sagittal (horizontal) and frontal (coronal) viewing planes used to show spatial overlap.

#### Clinical Profile and Motor Relevance of Connectivity Findings

Considering the clinical characteristics of our participant groups, people with Parkinson’s disease and HOA did not differ significantly in their Montreal Cognitive Assessment (MoCA) scores (people with Parkinson’s disease: 25.49 ± 3.8; HOA: 26.92 ± 2.9; *t(51)* = −1.93, *P* = 0.06) (Table 2), indicating that the connectivity differences we observed are unlikely to be driven by cognitive deficits. Instead, these differences appear more aligned with motor symptomatology. All people with Parkinson’s disease participants had elevated scores on the MDS-UPDRS, which directly indexes motor impairment (Table 2).

**Table 2.**
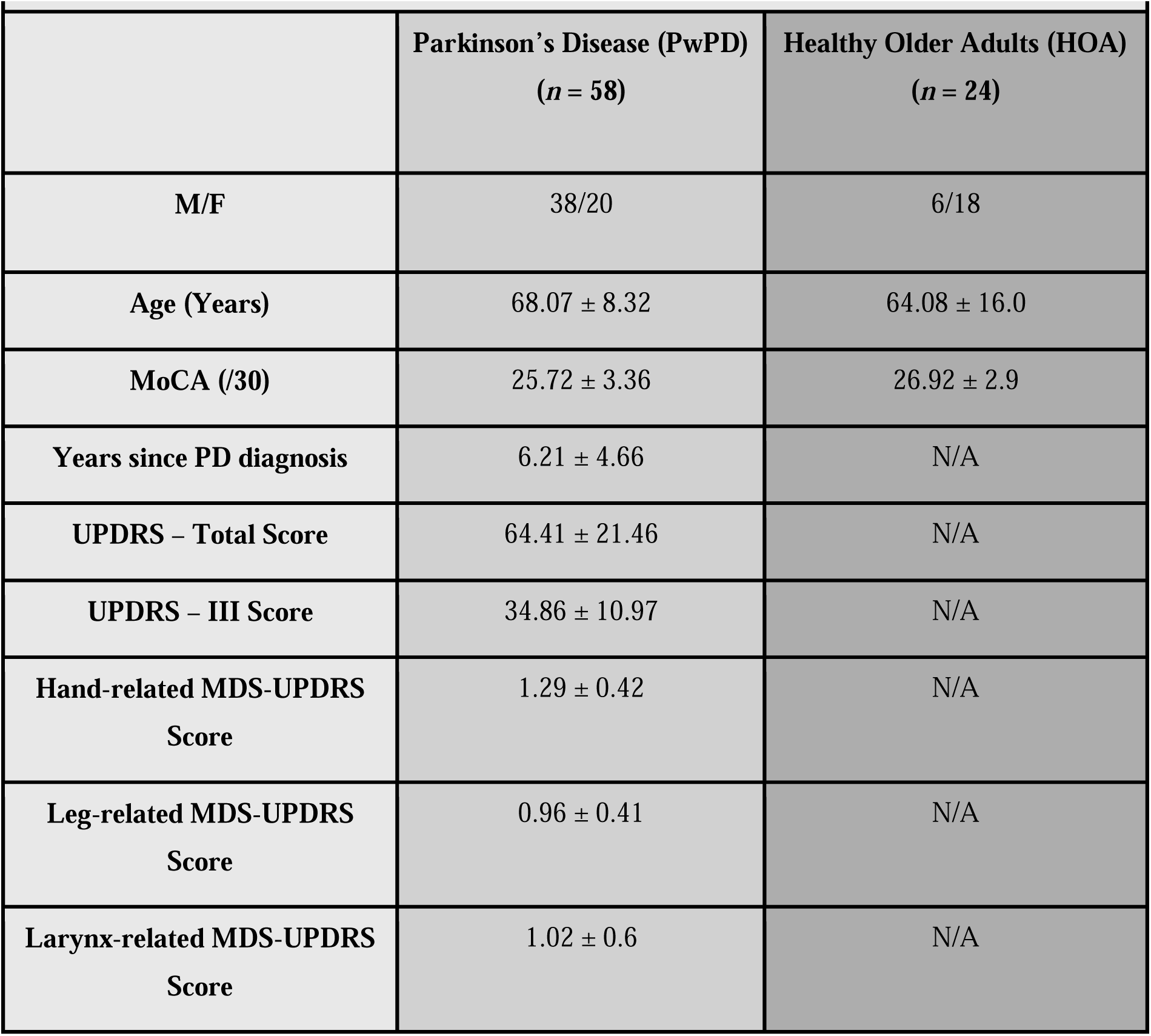
Participant demographics and clinical characteristics. This table presents demographic and clinical data (mean ± standard deviation) for both People with Parkinson’s Disease and HOA, including the number of male and female (M/F) participants, average age (in years), and mean Montreal Cognitive Assessment (MoCA) scores. For PwPD, additional data include the number of years since PD diagnosis, average Unified Parkinson’s Disease Rating Scale (UPDRS) Total Score, UPDRS-III (Motor Assessment) mean score, Hand-related UPDRS mean score, Leg-related UPDRS mean score, and larynx-related UPDRS mean score (Supplementary Tables 6, 7). N/A: Not Applicable.

| **Table 2. Participant demographics and clinical characteristics.** |  |  |
| --- | --- | --- |
|  | **Parkinson's Disease (PwPD)**<br>**(n = 58)** | **Healthy Older Adults (HOA)**<br>**(n = 24)** |
| **M/F** | 38/20 | 6/18 |
| **Age (Years)** | 68.07 ± 8.32 | 64.08 ± 16.0 |
| **MoCA (/30)** | 25.72 ± 3.36 | 26.92 ± 2.9 |
| **Years since PD diagnosis** | 6.21 ± 4.66 | N/A |
| **UPDRS – Total Score** | 64.41 ± 21.46 | N/A |
| **UPDRS – III Score** | 34.86 ± 10.97 | N/A |
| **Hand-related MDS-UPDRS Score** | 1.29 ± 0.42 | N/A |
| **Leg-related MDS-UPDRS Score** | 0.96 ± 0.41 | N/A |
| **Larynx-related MDS-UPDRS Score** | 1.02 ± 0.6 | N/A |

To link our rsfMRI findings more directly to symptom topography, we conducted an effector-specific MDS-UPDRS analysis. Using a predefined mapping of MDS-UPDRS-III items to leg-, hand-, and larynx-related motor functions (Supplementary Table 7), we calculated per-item mean severity scores for each effector category. Average scores indicated mild impairment across all categories (≈1/4 of the maximum possible score per item): Leg = 0.96 ± 0.41, Hand = 1.29 ± 0.42, and Larynx = 1.02 ± 0.6 (Supplementary Tables 8a-b). Despite this generally mild presentation, hand-related scores were modestly but significantly higher than those for the larynx or leg (Holm-adjusted Wilcoxon tests: Hand > Larynx, *p_adj* = 0.000252; Hand > Leg, *p_adj* = 0.000005), whereas larynx and leg scores did not differ significantly (*p_adj* = 0.63) (Supplementary Table 8c).

In our cohort, this pattern suggests that hand-related deficits may contribute disproportionately to connectivity findings in M1 subregions with strong representation of the hand area. For example, in IC54, impairments in hand tasks, more so than in leg, could underlie the observed increases in connectivity with the IFG (Figures 1, 3). Similarly, for IC66, hand-related impairment may help explain the heightened connectivity with the CB, more so than larynx-related deficits (Figures 1, 3). Nonetheless, because all three effectors showed mild impairment, the connectivity differences we identified could also reflect the combined influence of deficits across multiple motor systems. Together, these findings suggest that the altered connectivity patterns in people with Parkinson’s disease may reflect targeted reorganization in motor circuits affected by the disease, aligning both with anatomical specificity in M1 and with clinical measures of motor function.

## Discussion

In this study, we aimed to compare functional connectivity patterns in people with Parkinson’s disease and HOA, focusing on brain regions associated with IG and EG movement pathways. Our results revealed patterns of both increased and decreased functional connectivity across key motor and other areas. Specifically, the PreCG/M1 areas we tested showed only patterns of higher connectivity in people with Parkinson’s disease, when compared to HOA, specifically with cerebellar areas. Other cortical areas, such as most PoCG/S1 components also showed higher connectivity with cerebellar areas, while other areas, such as the insula and the caudate showed lower connectivity with cerebellar areas. Some areas, like the caudate, showed patterns of both higher connectivity (e.g., with the STG), and lower connectivity (e.g., with the SMFG). This divergence in connectivity trends underscores the complexity of connectivity alterations in people with Parkinson’s disease, highlighting region-specific variability rather than a uniform trend. It may also reflect underlying compensatory mechanisms, such as increased PreCG/M1-CB coupling, alongside disrupted or maladaptive patterns, such as caudate-SMFG interactions.

Our motor effector-specific analysis of the observed patterns in M1 revealed that some identified differences might be driven by effector-specific control, such as increased connectivity between leg, hand, and laryngeal areas with different cerebellar areas. This could support the idea that functional reorganization in people with Parkinson’s disease may be shaped not only by anatomical location within M1 but also by the specific motor effectors subserved by each region, with distinct effector-related circuits showing unique connectivity alterations. Analysis of MDS-UPDRS scores in our people with Parkinson’s disease cohort showed mild impairment across all effectors but significantly greater hand-related deficits compared to leg or larynx, suggesting that connectivity increases in hand-dominant M1 components may be partly driven by these disproportionate impairments. Our findings suggest that some of our identified differences may reflect compensatory mechanisms, with some of them tuned to effector-specific impairments.

### Comparison with Previous Studies

Several prior resting-state fMRI studies have described altered connectivity in people with Parkinson’s disease; however, they often used broad anatomical terminology (e.g., sensorimotor network, default mode network, auditory network)^53,54^ and diverse analytic frameworks (e.g., nodal graph theory measures or network efficiency),^55,56^ limiting direct pairwise comparisons with our study. Despite this, some parallels emerge. For instance, greater cerebellar connectivity in people with Parkinson’s disease has been reported,^26,56,57^ though often without specification of the cerebellar subregions or targets. Our results specify increased connectivity between specific subareas of the PreCG/M1(e.g., dorsomerdial, dorsolateral) and PoCG/S1 (e.g., dorsal, dorsomedial, dorsolateral) cortices with specific cerebellar subregions including Crus II, Lobule VIIIa, and VIIIb. A closer look at Kaut et al.,^26^ for example, shows that increased cerebellar connectivity in Parkinson’s disease-fallers (vs. HOA) involved different targets (e.g., vermis) within a distinct group comparison framework (Parkinson’s disease-non-fallers, Parkinson’s disease-fallers, and HOA), a relationship that did not hold up for the Parkinson’s disease-non-fallers vs. HOA comparison. These distinctions underscore the anatomical precision afforded by our approach and highlight the added value of high-resolution IC mapping for clarifying disease-related connectivity changes.

Where specific pairwise relationships were reported in other studies, some findings converge with ours. Hacker et al.^58^ noted reduced connectivity between the striatum (without specifying which striatal area) and cerebellum in people with Parkinson’s disease, paralleling our observed decrease between the striatal area (caudate) and CB Crus II. Müller-Oehring et al.^24^ identified the caudate as a region with altered connectivity in people with Parkinson’s disease, as we did, but the target regions reported were different than ours. Specifically, Müller-Oehring et al.^24^ found increased connectivity with the thalamus and insula, and decreased connectivity with motor and premotor regions, while our caudate-connectivity findings focused on increased caudate-STG connectivity, and decreased caudate-SMFG and caudate-cerebellar connectivity. Considering these comparisons, our approach offers fine-grained anatomical resolution and allows for translation between other studies’ findings, especially those reporting specific pairwise relationships and trends of connectivity (increased vs. decreased).

### Interpretation Through the Lens of IG/EG Movement Pathways

Although our study examined functional connectivity and most studies on IG/EG movement assess activation patterns, we attempt here to bridge these findings to provide insights into the functional significance of our observed connectivity differences. For instance, the cerebellum is one prominent brain region with altered connectivity in our study. Prior fMRI studies showed increased cerebellar activation during both IG and EG movements in healthy adults,^59–62^ while in people with Parkinson’s disease, cerebellar activation is more prominent during IG movements.^20,21,27,63,64^ Although our study assessed functional connectivity and not task-based activation, our observation of increased connectivity in people with Parkinson’s disease between cerebellum and other key areas, importantly the M1, may similarly reflect a compensatory mechanism specifically for IG movements. IG movements are more impaired in people with Parkinson’s disease, largely due to dysfunction in basal ganglia circuits,^28–30^ and increased connectivity with the cerebellum, a region critical for motor coordination and error correction,^29,30,56,65^ may suggest that it supports or supplements impaired basal ganglia pathways during IG movement execution. This hypothesis is further supported by findings that increased cerebellar connectivity is observed in people with Parkinson’s disease with tremor^66^ and those who fall,^26^ while increased M1 nodal connectivity has also been reported in people with Parkinson’s disease with tremor.^67^

Additionally, the motor regions showing altered connectivity in our study corresponded to effector-specific zones within M1, namely, leg-, hand-, and larynx-related areas. This effector-specific resolution allowed us to interpret differential cerebellar-M1 connectivity in light of muscle group-specific impairments. Many of the studies reviewed (Supplementary Table 1) used IG/EG tasks involving hand and leg movements, aligning with our findings of increased connectivity between cerebellar subregions and M1 zones associated with these effectors. Another effector that showed higher connectivity between M1 and the cerebellum is the larynx. Although this effector is not typically studied in the context of IG vs. EG movement distinctions, we hypothesize that our findings may correspond to known impairments in vocal motor control in people with Parkinson’s disease,^68^ which manifest as a range of speech motor deficits. Importantly, our cohort did include clinical assessments relevant to speech: the larynx-related MDS-UPDRS items (covering speech, saliva/drooling, chewing/swallowing, and eating tasks) demonstrated mild but measurable impairment. While the mean severity of larynx-related deficits was like that of leg-related deficits and lower than that of hand-related deficits, these scores nevertheless indicate the presence of vocal-oropharyngeal motor dysfunction in this sample. Considering that people with Parkinson’s disease show more deficits in their speech-related vocal motor control compared to the control needed for vocal behaviors such as laughing or yawning,^12^ our findings may reflect differences in control of voluntary (speech) versus more spontaneous vocalizations.

### Clinical Implications and Intervention Relevance

Our results reinforce the notion that IG movement pathways are more impaired in people with Parkinson’s disease, which could guide therapeutic strategies. Interventions leveraging EG movement cues, such as rhythmic cueing, tango, and tai chi, may harness relatively preserved networks to compensate for IG deficits.^69–71^ For instance, Argentine tango emphasizes rhythmic, externally guided motion, and has been shown to improve gait, mobility, and cognitive function in people with Parkinson’s disease compared to control groups that did not receive tango-based interventions.^72^ Our connectivity findings suggest a neural basis for these intervention effects, providing a framework for identifying relevant seed regions to measure brain connectivity changes pre- and post-dance-based intervention in future studies.

### Limitations

This study has several limitations. Our interpretive framework is grounded in prior literature identifying specific brain regions as being associated with IG, EG, or both movement pathways. Accordingly, we focused on a set of predefined regions of interest (ROIs) rather than conducting a whole-brain analysis. As a result, our findings are limited to the functional connectivity patterns observed within the selected ROIs and their corresponding Neuromark ICs, which may not capture thus far unidentified regions relevant to IG, EG, or combined movement control. While the use of NeuroMark components offers important advantages, such as standardized and replicable IC definitions across studies, the predefined nature of these components may overlook finer-scale dynamics involved in movement generation. The same caveat applies to our use of M1 effector masks, which were derived from prior analyses. Another limitation is that movement-related brain regions are not exclusively tied to either IG or EG pathways; their involvement can vary depending on the specific task being performed. Consequently, while our framework provides a useful lens, its reliance on binary classifications may not reflect the full complexity of the underlying neural architecture. Because our approach builds on existing literature, our conclusions are also shaped by the methodological assumptions and rigor of those prior studies. Any inconsistencies or limitations in that body of work may influence our interpretations. Despite these constraints, we are confident that the connectivity alterations identified here represent robust, anatomically meaningful, and replicable differences between people with Parkinson’s disease and HOA, even if the precise functional relevance to IG or EG pathways remains to be clarified.

## Data availability

The authors confirm that, for approved reasons, some access restrictions apply to the data underlying the findings. Public data deposition is not ethical or legal and would compromise patient privacy. Data are, in part, housed in the VINCI Veterans Affairs (VA) Informatics and Computing Infrastructure (VINCI), a secure, virtual computing environment developed through a partnership between the VA Office of Information Technology (OI&T) and the Veterans Health Administration’s Office of Research and Development (VHA ORD). Data are also stored on a VA secured server under a Data Usage Agreement with Emory University. VINCI is a partner with the Corporate Data Warehouse (CDW) and hosts all data available through CDW. VHA National Data Services (NDS) authorizes research access to patient data. Data are only available for researchers who meet the criteria for access to confidential data. Contact. Detailed analysis and results files of aggregated data can be found at https://github.com/nehabajaj101/PwPD-vs.-HOA/tree/main.

## Supporting information

Supplementary References

## Acknowledgements

We thank the volunteers and participants for their time and effort devoted to this study.

